# First Detection of Recombinant Enterovirus G Carrying a Torovirus Papain-like cysteine protease Gene from India

**DOI:** 10.1101/2024.05.01.591987

**Authors:** PM Sawant, M Lavania, A Kulkarni

**Author notes:** **Correspondence to:** Sawant P.M., Enteric Viruses Group, ICMR - National Institute of Virology, 20-A, Ambedkar Road, PO Box No 11, Pune 411 001, India.

## Abstract

Enterovirus G (EV-G) comprising 20 genotypes (G1-G20) are associated with in a various disease conditions in pig populations. The recombination between homologous EV-G is a driving force behind evolution but the recombination with virus from heterologous family is rare event. Presently, two types of recombinant EV-G strains containing torovirus papain like cysteine protease (PLCP) at 2C-3A junction (Type I) and in place of structural proteins (Type II) are reported. The present study conducted to reveal RNA virome in diarrheic piglet’s faeces retrieved genome coding for complete polyprotein of EV-G1. The detected EV-G1 strain represents first type I recombinant from India. The polyprotein of strain exhibiting 97% amino acid identity was closely related with Chinese strain. Furthermore, the EV-G1-PLCP strain was found to be originated from South Korean, Chinese, and Japanese strains with breakpoints at nucleotide (nt) position 1323 and 4725. Genetic and phylogenetic analysis of VP1 and PLCP of study strain demonstrated its closeness to EV-G1-PLCP strains from South Korea KOR/KNU-1811/2018/G1-PLCP. The present study advances the knowledge about genetic evolution of EV-G from gaining virulence gene from different virus family.

---

Enteroviruses, family *Picornaviridae*, infect human (EV-A to D), non-human primates (EV-A, EV-B, EV-D, EV-H & EV-J), bovine (EV-F & EV-E), and swine (EV-G), are small, non-enveloped viruses containing positive sense single stranded RNA genome. The EV genome possess a single open reading frame (ORF) flanked by 5’ untranslated region (UTR) and 3’UTR, and poly A tail. The ORF translates into single polyprotein, which is then posttranslationally processed to four capsid proteins (VP1-VP4) and seven functional proteins (2A-2C and 3A-3D). The untranslated regions at 5’UTR and 3’UTR of genome are essential for RNA replication (Zell et al., 2017).

The EV-G is frequently detected from pigs worldwide, clinical relevance of virus apart from pyrexia, skin lesion, flaccid paralysis has not been investigated. The EV-G currently includes 20 genotypes, which undergo intragenotypic and intergenotypic recombination or cross-order recombination between EV-G (Picornavirales), and torovirus (family *Coronaviridae* and order *Nidovirales)* (Muslin et al., 2015). Recently, two types of EV-G variants arising from recombination of G1, G2, G8, G10, G12, G17 genotypes and papain-like cysteine protease sequence (PLCP) from torovirus are reported (Li et al., 2023). The type 1 recombinant have insertion sequence at 2C-3A junction and detected from fecal samples of porcine diarrhea cases in USA, Belgium, Germany, South Korea (Bunke et al., 2018; Knutson et al., 2017; Shang et al., 2017; Conceição-Neto et al., 2017; Lee & Lee, 2019), while from asymptomatic and diarrheic pigs in Japan (Tsuchiaka et al., 2018). The recombinant type 2 viruses containing torovirus PLCP gene flanked by long unique genes 1 and unique region gene 2 in the place of viral structural genes has been detected in Chinese and Japanese pigs (Wang et al., 2018; Imai et al., 2023). Another novel type 2 recombinant replacing viral structural protein genes by PLCP gene with upstream (without homology with GenBank deposited protein) and downstream (homology with a baculoviral inhibitor of apoptosis repeat superfamily) sequences was detected in Japanese pig faeces (Imai et al., 2019). The PLCP gene present in ORF1 of torovirus serves as protease to cleave viral ORF1a and ORF1b polyproteins to nonstructural proteins (Mielech et al., 2014). Additionally, it can influence pathogenicity by acting as innate immunity antagonist through deubiquitinating and deISGylating (interferon stimulating gene 15-removing) enzyme activity (Shang et al., 2017).

In the present study, next generation sequencing (NGS) performed on rotavirus group A (RVA) positive samples from a single farm led to detection of a recombinant EV-G detected in diarrheic piglet that contained 223 AA insertion of ToV-PLCP and characterized its full genome to explore new genetic features (Sawant et al., 2023). The Indian recombinant type 1 EV-G1 possesses single ORF of 7161 nucleotide encoding 2387 amino acid (AA) polyprotein, which started at translational start site (ATG) at 5’ end and ended at TAA after 3D^pol^ gene. The gene order in type 1 recombinant EV-G1 detected in present study was: 5’UTR, VP4, VP2, VP3, VP1, 2A, 2B, 2C, PLCP, 3A, 3B, 3C, 3D, 3’UTR comprised 69, 246, 238, 282, 150, 99, 329, 223, 89, 22, 183, 461 AA respectively.

The potential cleavage site predicted after alignment with reference genomes was mainly Gln(Q)/Gly(G) which is cleaved by 3C^pro^ or 3CD^pro^, except for cleavage sites at junctions of VP4/VP2, VP3/VP1, VP1/2A, 2B/2C, which were Lys (K)/Ser (S), Gln (Q)/Ala (A), Thr (T)/Gly (G), Gln (Q)/Asn (N). After alignment of recombinant EV-G sequences, the 3C protease cleavage site predicted as ALFQ/GPPA and AVFQ/GPPA flanked 5’ and 3’ prime sites of PLCP insert avoiding the disruption of proteolytic cleavage of polyprotein (Fig 1). In India, EV-G has been reported but torovirus have not been detected from pigs. However, detection of recombinant EV-G1-PLCP but not porcine torovirus indicates that recombination might have took place in the past and circulation of torovirus in Indian pig population.

**Fig 1:**
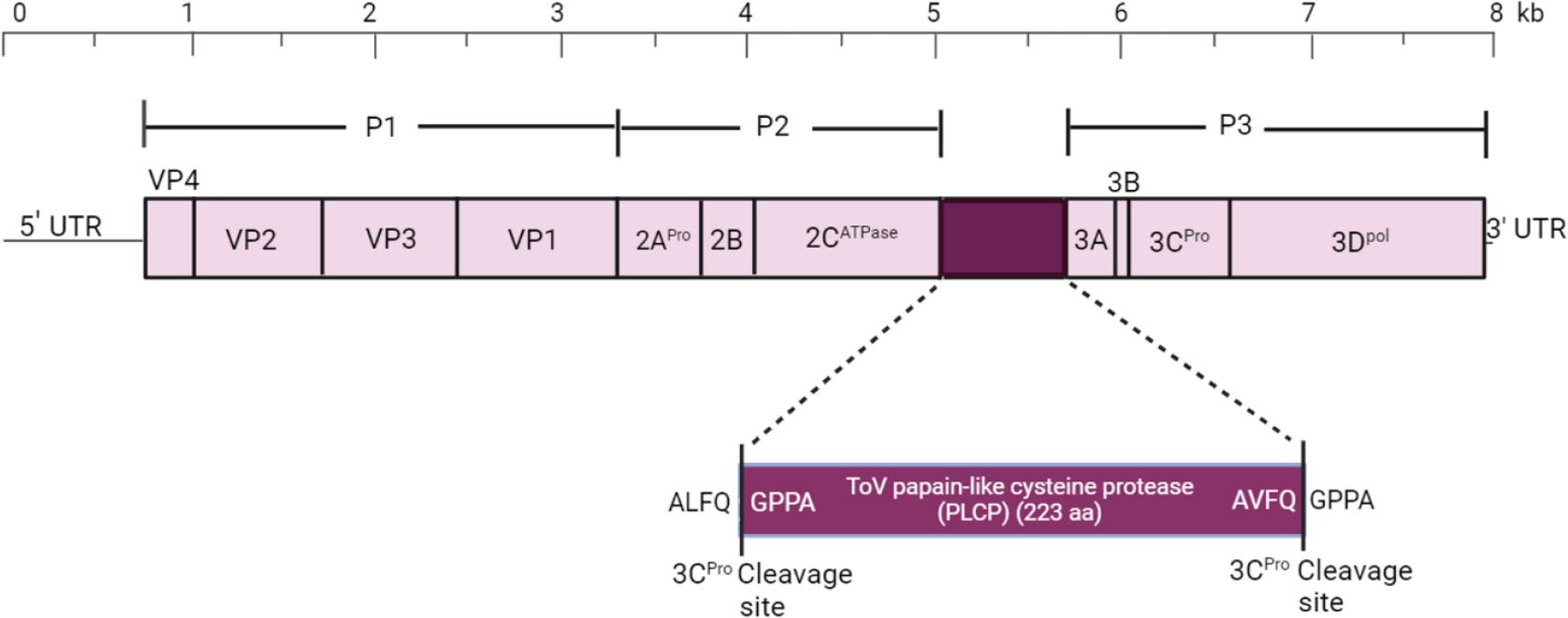
Schematic presentation of enterovirus G1-PLCP genome: The virus codes for single polyprotein (divided into P1, P2, and P3 regions) which cleaved into 11 mature proteins. The purple box at 2C-2A shows torovirus papain like cysteine protease (PLCP). The box also shows 5’ and 3’ boundaries of 3C protease cleavage sites.

The confirmation of insert as part of original viral genome and not assembled from coinfection with other unrelated virus was performed by assessing overlapping read coverage across boundaries. The amino acid insert sequence boundaries were found by running a translated BLAST search (tBLASTn) against a previously assembled nucleotide reference contig (contig00001-NIV1740787) using standalone BLAST version 2.8.1. The number of reads that overlap the insert region were determined by performing a BLASTn search using standalone BLAST. Using a module in FASTX Toolkit 0.0.14, the reads that were originally in fastq format were translated to FASTA format and then used for BLAST search. The insert amino acid sequence which spans 223 amino acids long, starts at 4582 nt and terminates at 5250 nt of the reference contig. The number of reads that span the exact insert region are 348 R1 reads and 352 R2 reads. 227 R1 reads and 187 R2 reads were mapped at the borders aligning with at least one base of the insert. The range of 149 nt prior to and 149 nt after the boundary was taken into consideration in order to calculate the read abundance at the boundaries. Similarly, BLASTn search was used to map the read abundance at the gene boundaries for the other genes. The General feature format file (GFF file) that describes the features of each gene was used to determine the borders for each gene.

The unique insertion sequence found in EV-G genome at 2C-3A junction was subjected to BLASTn and found to be homologous to recently detected recombinant enteroviruses from USA, Japan and Belgium (Bunke et al., 2018; Knutson et al., 2017; Shang et al., 2017; Conceição-Neto et al., 2017; Lee & Lee, 2019). However, the length of PLCP insert varied from different geographical regions (198-211 AA in Korean strains, 197-217 AA in Japanese strains, 194-223 AA USA strains, 217 AA Chinese strains). Next, sequence identity and phylogenetic relationship was identified between study strain, recombinant EV-G1-PLCP full genomes and torovirus sequences available in the GenBank (Fig 2). The PLCP of study strain showed 81.6-89.6 % AA identity with G1, G2, G10, G17 strains, showing maximum identity with KNU-1811 strain (G1-PLCP). Similar to previous observations the PLCP insert of EV-G study strain shared greater amino acid identity with reported recombinant EV-G than torovirus, indicating that the recombination event has happened in the past (Fig 3). Phylogenetic analysis shows that complete polyprotein and VP1 gene of novel study strain shared maximum identity in the range of 94-97% and 94-95.4% AA identity to EV-G-PLCP (G1 genotype) detected in pigs from the China, South Korea, Japan, and USA. The recent study covering wider areas of India suggested that EV-G (G1, G6, G8 and G9) types are in circulation throughout the country. However, they could not find recombinant EV-G as smaller portion of genome was targeted for characterization jeopardizing the finding of the origin of recombinant study strain.

**Fig 2:**
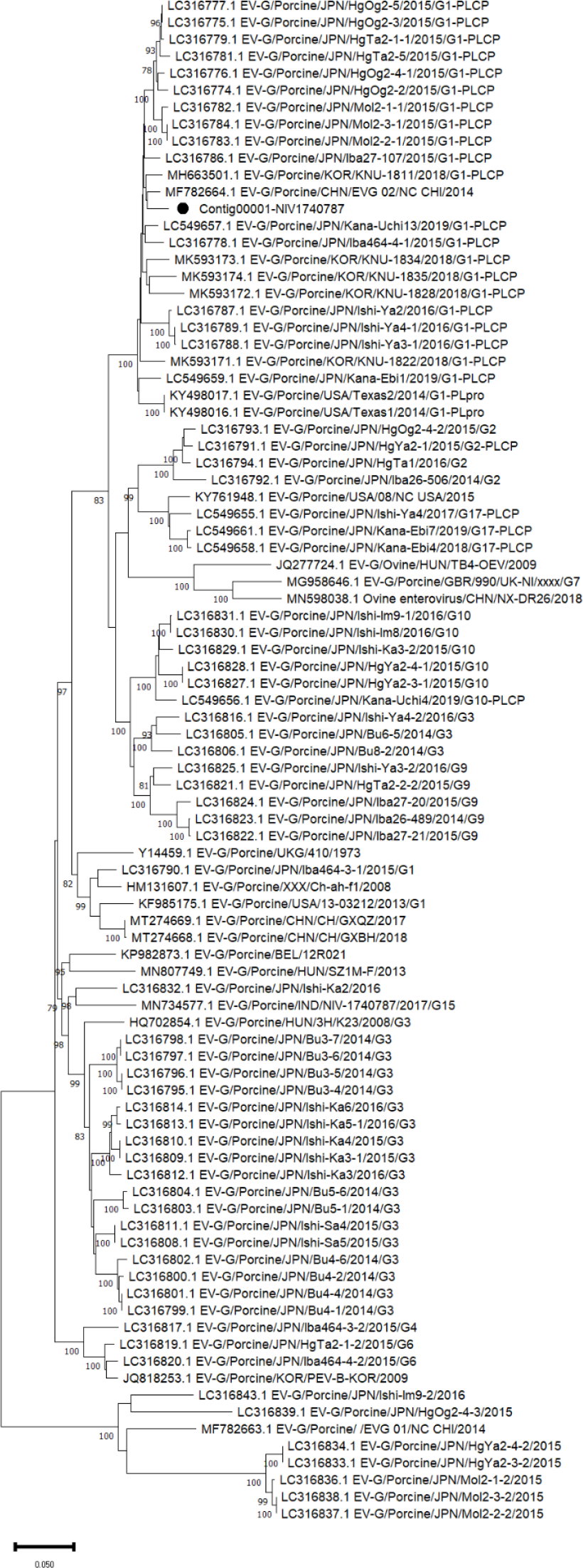
Phylogenetic tree of enterovirus G polyprotein: Multiple sequence alignment was performed with clustal W and the Neighbor Joining method with 1000 bootstrap replicates (branches shows values >75%) was used to construct the tree. The study strain is marked with black circle.

**Fig 3:**
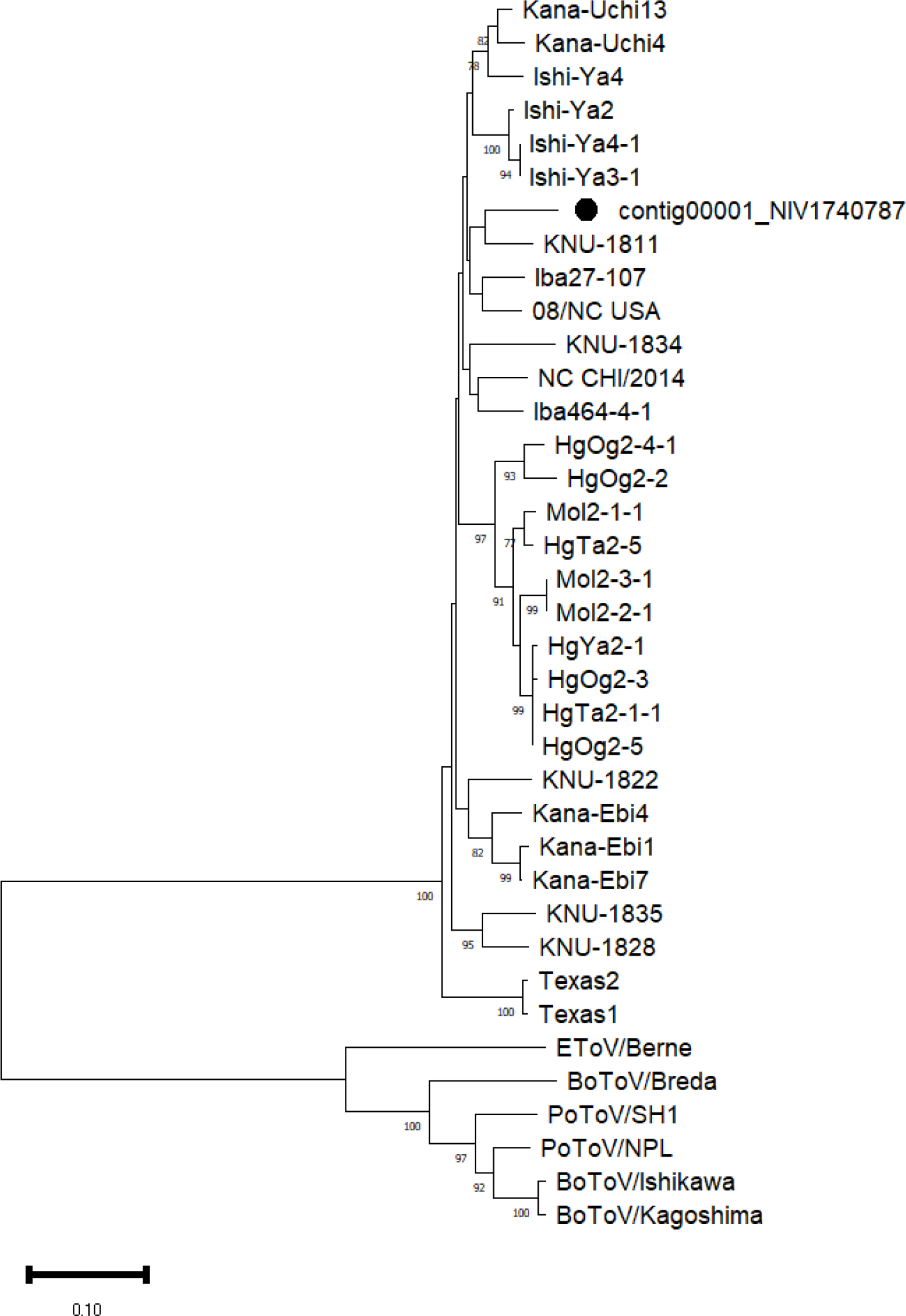
Phylogenetic tree of papain like proteases of recombinant enterovirus G and torovirus: The tree was constructed using neighbor Joining method with 1000 bootstrap replicates (percent values on branches). The study strain is indicated with black solid circle.

The enteroviruses evolve rapidly by high point mutations and frequent recombination (Imai et al., 2019). Genome recombination analysis in boot scan suggested that EV-G strain (Contig00001-NIV1740787) evolved from parents from South Korea (KOR/KNU-1835/2018/G1-PLCP), China (EV-G 02/NC_CHI/2014), and Japan (EV-G/Porcine/JPN/Bu8-2/2014/G3), with a breakpoint observed at nucleotide position 1323 and 4725 (Fig 4). The detection of recombinant virus indicates that the pig trades and faeco-oral transmission are the facilitators of recombination between the viral strains from different geographical locations. The main constraint of recombination studies is the unavailability of full genome sequences, as EV-G typing is performed based on partial VP1 sequencing. Accumulation of more full genomes will reflect the diversity of circulating strains and in future may yield different findings and conclusions.

**Fig 4:**
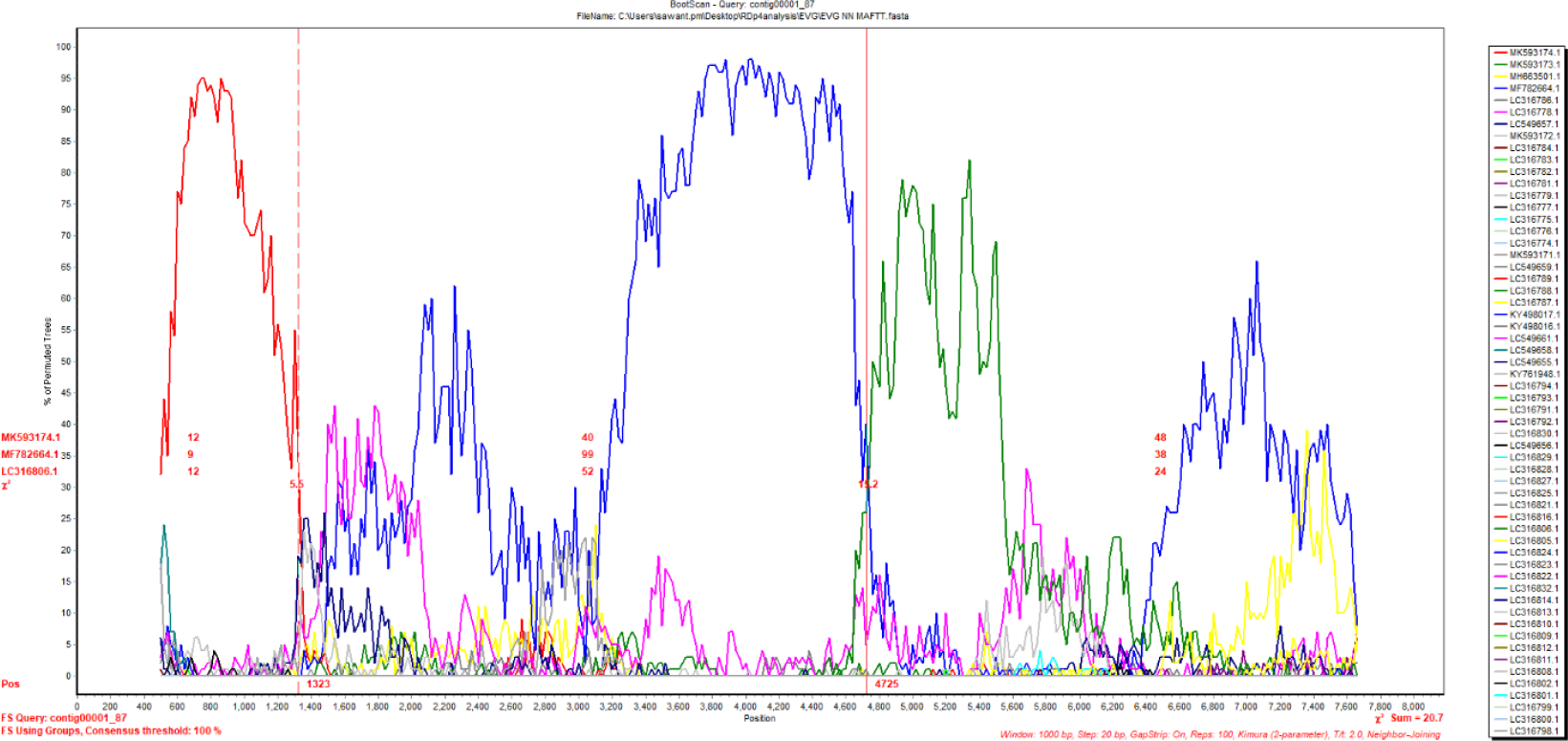
The bootscan plot exhibiting recombination events between Indian EV-G1-PLCP and parent strains: KOR/KNU-1835/2018/G1-PLCP strain (MK593174.1), EV-G 02/NC_CHI/2014 strain (MF782664.1), and EV-G/Porcine/JPN/Bu8-2/2014/G3 strain (LC316806.1). The analysis was performed by neighbour-joining algorithm with sliding window size of 800bp, step size of 10bp, 100 bootstrap replicates, and gap-stripped alignments.

Next generation sequencing has detected coinfections with multiple porcine enteric viruses in faeces, including rotavirus, astrovirus, enterovirus, circovirus, picobirnavirus, teschovirus, sapelovirus, kobuvirus, posavirus, pasivirus (Capai et al. 2022). Accordingly, in this study EV-G detected along with viral coinfections rotavirus, astrovirus, picornavirus, picobirnavirus (Sawant et al., 2023). The role of diverse viruses in diarrheic and healthy piglets is unknown. It seems that viruses are part of healthy intestinal virome and few viruses under some conditions are able to cause disease, one of them is infection with virulent viruses like rotavirus, transmissible gastroenteritis virus and porcine epidemic diarrheal virus. Another way the virulence of EV-G is influenced by the acquisition of foreign innate immune antagonist, torovirus PLCP. Additionally, experimental proven pathogenic potential of G1-PLCP genotype (Mi et al., 2021), the capacity of EV-G-PLCP to infect cell lines from diverse origins (Li et al., 2023) and non-homologous recombination between EV-G (picornavirus) and torovirus (coronavirus) highlights their potential to cross species barriers.

In conclusion, the present study first time detected recombinant EV-G genotype G1 from *Picornavirales* carrying viral gene of torovirus belonging to *Nidovirales* from India. In future, the studies should investigate the role of PLCP insertion on the association of EV-G with disease in Indian pigs.

## Author contribution

Conceptualization, PMS; Methodology, PMS, ML, AK; Data analysis, PMS, AK; Manuscript writing, PMS; AK, ML.

